# Rapid Online Assessment of Reading Ability

**DOI:** 10.1101/2020.07.30.229658

**Authors:** Jason D. Yeatman, Kenny An Tang, Patrick M. Donnelly, Maya Yablonski, Mahalakshmi Ramamurthy, Iliana I. Karipidis, Sendy Caffarra, Megumi E. Takada, Klint Kanopka, Michal Ben-Shachar, Benjamin W. Domingue

## Abstract

An accurate model of the factors that contribute to individual differences in reading ability depends on data collection in large, diverse and representative samples of research participants. However, that is rarely feasible due to the constraints imposed by standardized measures of reading ability which require test administration by trained clinicians or researchers. Here we explore whether a simple, two-alternative forced choice, time limited lexical decision task (LDT), self-delivered through the web-browser, can serve as an accurate and reliable measure of reading ability. We found that performance on the LDT is highly correlated with scores on standardized measures of reading ability such as the Woodcock-Johnson Letter Word Identification test (r = 0.91, disattenuated r = 0.94). Importantly, the LDT reading ability measure is highly reliable (r = 0.97). After optimizing the list of words and pseudowords based on item response theory, we found that a short experiment with 76 trials (2-3 minutes) provides a reliable (r = 0.95) measure of reading ability. Thus, the self-administered, Rapid Online Assessment of Reading ability (ROAR) developed here overcomes the constraints of resource-intensive, in-person reading assessment, and provides an efficient and automated tool for effective online research into the mechanisms of reading (dis)ability.

---

Why is learning to read almost effortless for some children and a continuous struggle for others? Understanding the mechanisms that underlie individual differences in reading abilities, and contribute to reading disabilities, is an important scientific challenge with far reaching practical implications. There is near consensus on the importance of foundational skills like phonological awareness for reading development ^1–3^. However, there are a myriad of other sensory, cognitive and linguistic abilities whose link to reading is fiercely debated ^4–11^. For example, hundreds of papers, published over the course of four decades, have debated the causal link between deficits in the magnocellular visual pathway and developmental dyslexia ^6,12-18^. Similarly contentious is the link between rapid auditory processing and developmental dyslexia ^19–23^. More recently, the role of various general learning mechanisms (e.g., statistical learning, sensory adaptation) in reading development has become a topic of great interest ^24–28^.

One reason these debates persist is that most findings reflect data from relatively small samples collected in a single laboratory and, therefore, findings are not necessarily representative of the population ^29–31^. There are two major factors that limit the feasibility of collecting data in large, diverse, representative samples. First, experiments measuring specific sensory and cognitive processes typically depend on software installed on computers in a laboratory. Second, standardized measures of reading ability typically require a trained test administrator to administer the test to each research participant. These two factors: (a) make it time consuming and costly to recruit and test large samples, (b) prohibit including research subjects that are more than a short drive from a university, (c) bias samples towards university communities and (d) create major barriers for the inclusion of under-represented groups.

Over the past decade, advances in software packages for running experiments through the web-browser have catalyzed many areas of psychological research to pursue larger, more diverse and representative samples. For example, software packages such as jsPsych ^32,33^, Psychopy ^34^, Gorilla ^35^ and others, make it feasible to present stimuli and record behavioral responses in the web-browser with temporal precision that rivals many popular software platforms for collecting data in the laboratory ^36^. However, online experiments have yet to make major headway in the developmental and educational spheres, specifically in reading and dyslexia research. Standardized testing of reading ability still depends on individually administered tests. Thus, even though a researcher might be able to accurately measure processing speed or visual motion perception in thousands of subjects through the web-browser, they would still need to individually administer standardized reading assessments to each participant.

Our goal here was to develop an accurate, reliable, expedient and automated measure of single word reading ability that could be delivered through a web-browser. Specifically, we sought to design a web experiment to approximate scores on the Woodcock-Johnson Letter Word Identification (WJ-Word-ID) test ^37^. The WJ-Word-ID is one of the most widely used standardized measures of reading ability in research (and practice). Like other test batteries that contain tests of single word reading ability (e.g., The NIH Toolbox ^38^, Wide Range Achievement Test ^39^, Wechsler Individual Achievement Test ^40^), the WJ-Word-ID requires that an experimenter or clinician present each research participant with single printed words for them to read out loud. Participant responses are manually scored by the test administrator as correct or incorrect based on accepted rules of pronunciation. The words are organized in increasing difficulty such that the number of words that a participant correctly reads is also a measure of the difficulty of words that the participant can read. Due to excellent psychometric properties, and widely accepted validity, the Woodcock-Johnson and a compendium of similar tests are at the foundation of thousands of scientific studies of reading development. Unfortunately, tests like the Woodcock-Johnson are not yet amenable to automated administration through the web-browser because speech recognition algorithms are still imperfect, particularly for children pronouncing decontextualized words and pseudowords.

In considering a suitable task for a self-administered, browser-based measure of reading ability, the lexical decision task (LDT) is a good candidate for practical reasons. Unlike naming or reading aloud, the LDT can be: (a) scored automatically without reliance on (still imperfect) speech recognition algorithms, (b) completed in group (or public) settings such as a classroom and (c) administered quickly as each response only takes, at most, 1-2 seconds. The LDT has a rich history in the cognitive science literature as means to probe the cognitive processes underlying visual word recognition. It is broadly assumed that some of the same underlying cognitive processes are at play when participants make a decision during a two-alternative forced choice (2AFC) lexical decision as during other word recognition tasks (e.g. naming ^41,42^). However, there are also major differences between the processes at play during a lexical decision versus natural reading ^43,44^. For example reading aloud requires mapping orthography to a phonetic and motor outputs with many possible pronunciations (as opposed to just two alternatives in an LDT). Moreover, selection of the correct phoneme sequence to pronounce a word, or even the direct mapping from orthography to meaning (as is posited in some models ^45^), certainly doesn’t require an explicit judgement of lexicality. Thus, from a theoretical standpoint there is reason to be optimistic that carefully selected LDT stimuli will tap into the key mechanisms that are measured by other, validated measures of reading ability (e.g., WJ-Word-ID). However, concerns about face validity would need to be dispelled through extensive validation against standardized measures since there are also clear differences between lexical decisions and reading aloud.

There is some empirical support for LDT performance being linked to reading ability. For example, Martens and de Jong demonstrated that response times (RT) on the LDT differed between children with dyslexia compared to children with typical reading skills ^46^. Specifically, they found larger word-length and lexicality effects in children with dyslexia compared to children with typical reading skills. These effects seem to track reading ability, as opposed to being a specific marker of dyslexia: reading matched controls; i.e., younger children matched to the children with dyslexia in terms of raw reading scores showed similar word-length and lexicality effects as found in the older children with dyslexia. Similar to our goal here, Katz and colleagues examined the LDT in combination with a Naming Task, as predictors of standardized reading measures ^47^. They found that average response times on the LDT (when combined with the Naming Task) correlated with a variety of reading ability composite scores, including the Woodcock Johnson Basic Reading Skills ^47^. However, their study primarily looked at young adults (median age of 21.5 years), and focused on RT as the independent variable, since most participants were at ceiling in terms of accuracy. It is not clear how well these effects would generalize to children learning to read. Moreover, effect sizes were relatively small, suggesting that RT on the LDT may not be the best predictor of reading ability, or that the range of stimuli was not broad enough. More recently, Dirix, Brysbaert and Duyck (2019) examined the relationship between LDT RT and eye tracking measures during natural reading and came to the conclusion that RT varies substantially across contexts and is not a stable measure of word recognition ^44^. Taken together, there is some evidence that (a) LDT performance is related to reading ability but (b) RT does not have suitable psychometric properties to construct a standardized measure. However, to our knowledge, no previous study has examined LDT response accuracy as a measure of word recognition ability in the context of development.

Here we sought to develop an implementation of the LDT in the web-browser, along with a suitable list of words and pseudowords, to efficiently and accurately measure reading ability across a broad age range, spanning first grade through adulthood. Specifically, we explore whether correct/incorrect responses to words of varying difficulty levels can serve as an accurate and reliable measure of reading ability. We, first, report results from a study in which we developed, validated and optimized a browser-based LDT measure of reading ability (*Study 1*). In *Study 1* we employed a long list of words and pseudowords, spanning a broad range of lexical and orthographic properties. The goals of *Study 1* were to, first, assess the feasibility of using a browser-based LDT to estimate reading ability and, second, leverage the item level data in combination with item response theory (IRT) to construct a collection of short-form tests with optimal psychometric properties. We then report a second study (*Study 2*) validating the optimized measure from *Study 1* to ensure that scores are reliable and consistent with standardized in-person measures: (a) for young children (6-7 year-olds) and (b) across a broad sample spanning early childhood through adulthood. We release the Rapid Online Assessment of Reading (ROAR), and the associated data and code, for other researchers to use (https://github.com/yeatmanlab/ROAR-LDT-Public).

## RESULTS: STUDY 1

### Lexical decision task performance and reading ability

To examine whether response times (RT) recorded in the online LDT replicated classic findings in the literature we fit linear mixed effects models to log transformed RT data for all correct trials. Each model included random intercepts for each word and random intercepts and slopes for each subject. We examined six different models containing fixed effects of (a) lexical status (real word vs. pseudoword), (b) log transformed lexical frequency (real words only), (c) log transformed bigram frequency, (d) word length, (e) the interaction effect of word length and participant reading ability as measured with the WJ-Word-ID and (f) lexical status, bigram frequency, word length and their interactions. As expected, we found: (a) RT was longer for pseudowords compared to real words (**β** = 0.11, SE = 0.0098, t = −11.35) (similar to ^48^); (b) responses to real words were faster with increasing lexical frequency (**β** = −0.021, SE = 0.0022, t = −9.84) (similar to ^48–50^); (c) responses to pseudowords were slower with increasing bigram frequency (**β** = 0.035, SE = 0.0062, t = 5.65) (similar to ^48,51,52^) and responses to real word were not affected by bigram frequency (**β** = −0.0033, SE = 0.0079, t = −0.43); (d) responses to words and pseudowords were slower with increasing stimulus length (**β** = 0.031, SE = 0.0046, 3 % increase per letter, t = 6.6) (similar to ^48^); and (e) there was a significant length by reading ability interaction (**β** = −0.010, SE = 0.0036, t = 2.76) indicating that the effect of word length on reading speed diminishes as reading ability improves (similar to ^46,48^). When lexical status, bigram frequency, word length and their interactions were entered into a single model (f), all the main effects were significant; the only significant interaction was between lexical status and bigram frequency indicating that bigram frequency influenced RT for pseudo but not real words (**β** = −0.032, SE = 0.0057, t = −5.87).

Stimuli for the LDT spanned a broad range of lexical and orthographic properties (**Supplementary Figure 1**), with the goal of providing a sufficiently large difficulty range for detecting associations between accuracy on the LDT and reading ability measured with the WJ-Word-ID. Indeed, overall accuracy, calculated as the proportion of correct responses on the 500 LDT trials, was highly correlated with WJ-Word-ID (**Figure 1 left panel**, r = 0.91, p < 0.00001, disattenuated r = 0.94). Accuracy for pseudowords (250 trials) had a slightly higher correlation with WJ-Word-ID (r = 0.86, p < 0.00001) than did accuracy for real words (r = 0.74, p < 0.00001), and this difference in correlation approached significance based on William’s test (p= 0.08; **Figure 1 right panel**).

**Figure 1.**
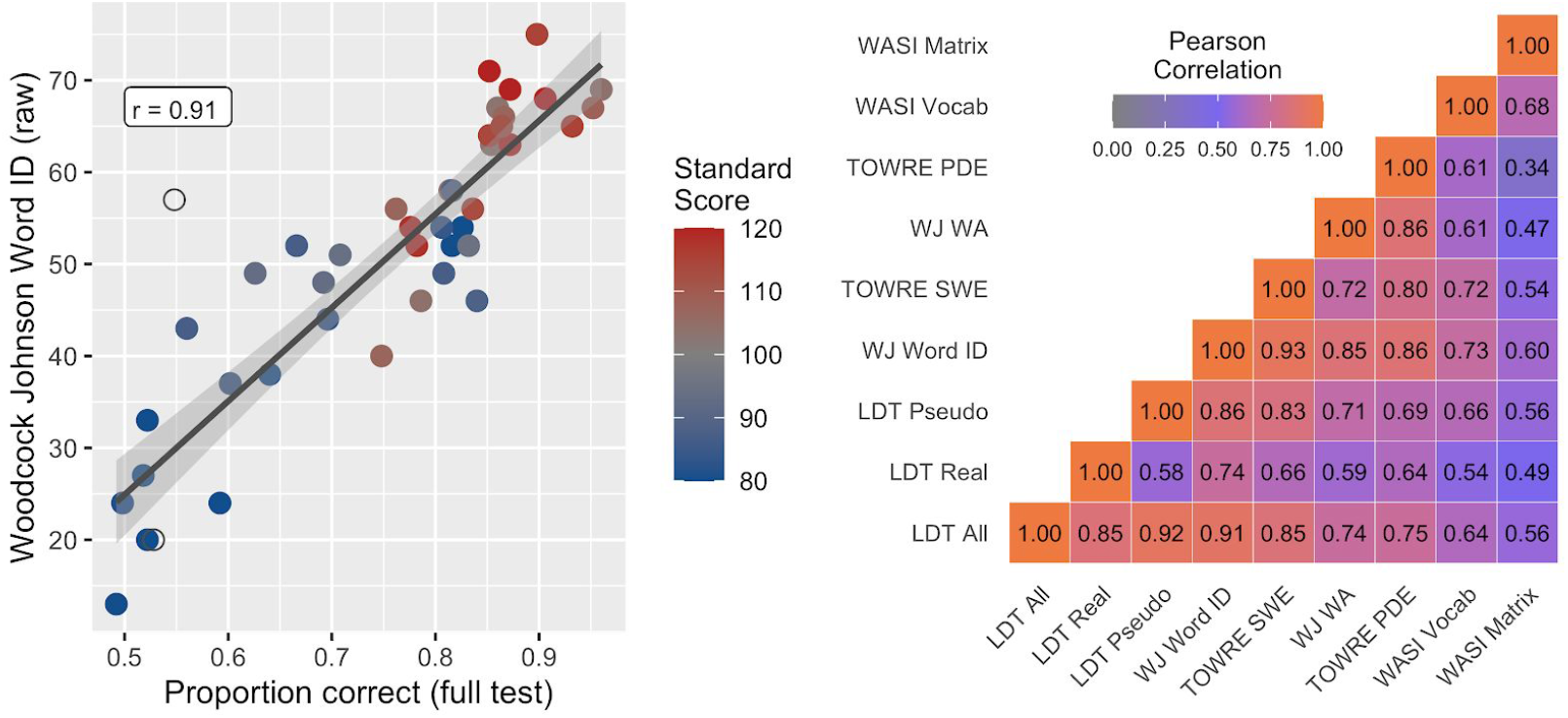
Accuracy on the lexical decision task predicts reading ability. (Left panel) Scatter plot showing each participants’ accuracy (proportion correct) on the 500 item LDT versus their raw score on the Woodcock-Johnson IV Letter Word Identification (WJ-Word-ID) test. Points are colored based on WJ-Word-ID standard scores, indicating that the relationship holds for struggling readers and those with dyslexia (dark blue), average readers (gray), above-average and exceptional readers (dark red). Open circles indicate outliers that were excluded from the analysis based on extremely fast response times on the LDT (see Methods). (Right panel) Correlation matrix between different measures of reading performance. The colormap represents the strength of correlation between each pair of variables. LDT All: overall accuracy on LDT; LDT Real: accuracy for real words; LDT Pseudo: accuracy for pseudowords. TOWRE SWE and PDE: Test of Word Reading Efficiency Sight Word Efficiency and Phonemic Decoding Efficiency; WJ WA: Woodcock Johnson Word Attack; WASI Vocab and Matrix: Wechsler Abbreviated Scales of Intelligence Vocabulary and Matrix Reasoning.

In contrast, median RT across correct responses in the LDT was not correlated with WJ-Word-ID (r = −0.06, p = 0.69). No correlations were found when median RT was calculated separately for words and pseudowords (r = −0.13, r = −0.08, respectively), even though RTs followed the expected patterns in other analyses (e.g., lexicality effects, see preceding paragraph). However, when the RT analysis was limited to the subset of the subjects who performed above 70% correct (N=28 with WJ-Word-ID scores), a significant negative correlation was found between RT and WJ-Word-ID (r = −0.53, p < 0.005). The correlation between WJ-Word-ID and median RT for real words (r = −0.53, p < 0.005) was similar to the correlation with median RT for pseudowords (r = −0.52, p < 0.005), and this difference was not significant according to William’s test (p = 0.79). Thus, response accuracy is a much stronger predictor of reading ability than RT (William’s test t =-5.72, p < 0.00005), and RT is only informative in relatively high performing subjects.

The impact of participant characteristics on the LDT-WJ correlation was examined through mediation and moderation analyses. The relation between LDT accuracy and WJ-Word-ID raw scores was not moderated by WJ-Word-ID standard scores, age, or number of months since testing (t = 0.71, p= 0.48; t = 0.12, p= 0.91; t = 1.24, p = 0.22). These results were further confirmed by a mediation analysis (CI [-0.001, 0.00], p= 0.69; CI [-0.001, 0.00], p= 0.87; CI [-0.0002, 0.00], p = 0.49). Together, these analyses suggest that LDT accuracy is a good approximation of standard reading measures for children with reading profiles ranging from severely impaired to exceptional, and ages between 6 and 18 years. It also suggests that LDT accurately predicts reading ability measured within the past 12 months.

Performance across different measures of reading ability is typically highly correlated. Indeed, different tests are frequently used interchangeably in different studies. For example, WJ-Word-ID scores and TOWRE-SWE scores are typically highly correlated (r=0.93 in our sample), even though the former is an untimed test while the latter is a timed test that combines real word reading accuracy and speed. In fact, it is common to use a threshold on either test to group subjects into dyslexic versus control groups in many research studies 21,53-55. The average Pearson correlation between the timed (TOWRE), untimed (WJ), real word (Word-ID and SWE) and pseudoword (Word Attack and PDE) standardized reading measures was r = 0.83 (range: r = 0.72 to r = 0.93; **Figure 1 right panel**). Given the correlation between LDT accuracy and WJ-Word-ID, it is not surprising that LDT accuracy was also significantly correlated with all the standardized reading measures (**Figure 1 right panel**). The correlations between LDT accuracy and Vocab (r = 0.64) and Matrix Reasoning (r = 0.56) were also nearly identical to the correlations between other reading measures and verbal abilities (r = 0.61 to 0.73) and reasoning abilities (r = 0.34 to 0.60, **Figure 1 right panel**). Thus, LDT response accuracy seems a suitable measure of reading ability as it behaves in a similar manner to the other established, standardized reading measures.

### Item response theory analysis of responses to individual words and pseudowords

Next we turn to the challenge of constructing a LDT with suitable psychometric properties for applicability in a research setting. The word list used in Study 1 was constructed to broadly sample different types of words and pseudowords. In doing so, our goal was to use item response theory (IRT) to select the optimal subset of stimuli that measure reading ability reliably and efficiently across the full continuum of reading abilities. Our approach was to remove items that were: (a) not correlated with overall test performance and WJ-Word-ID performance, (b) not well fit by the Rasch model and (c) examine item disrcimination based on the two parameter logistic (2PL) model. We then constructed an optimized set of three short stimulus lists, matched in terms of difficulty.

### Step 1: Remove words that are not predictive of overall performance

As a first step, we computed the correlations between item responses, LDT accuracy calculated on the full 500 item test, and WJ-Word-ID scores. Performance on some stimuli, like the pseudoword “insows”, was highly correlated with overall test performance (r = 0.61, p < 0.00001) and WJ-Word-ID score (r = 0.60, p = 0.000035). This means that a participant’s response to that pseudoword was highly informative of their reading ability. Other stimuli, like the pseudoword “timelly”, and the real words “napery” and “kind”, provided no information about overall test performance, as indicated by correlations near (or below) zero. We removed 71 items that did not surpass a lenient threshold of r = 0.l0. **Figure 2** shows words that were rejected (red) and retained (blue) based on this criterion.

**Figure 2.**
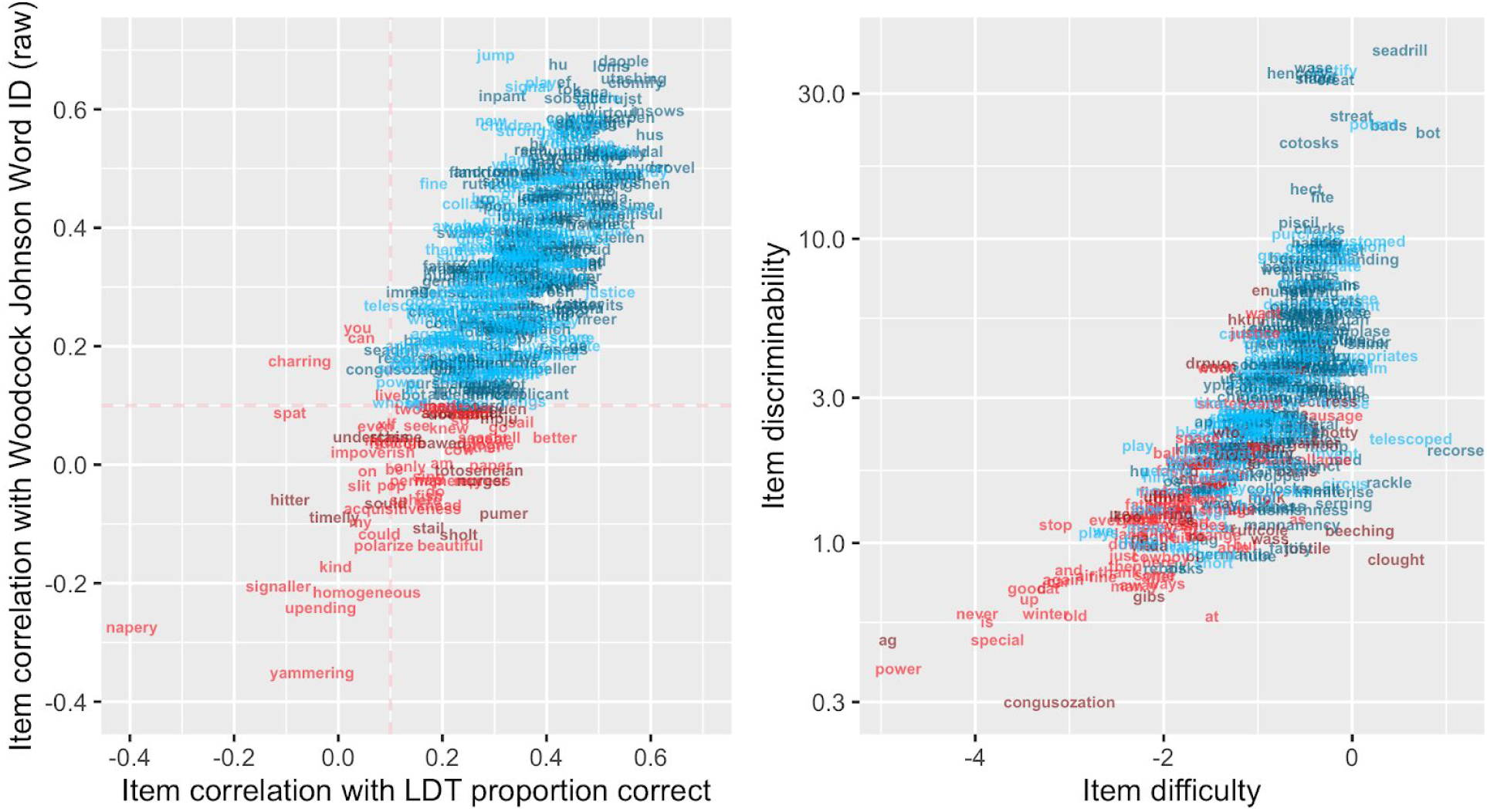
Item analysis. (Left panel) For each stimulus, we plot the correlation between performance on this item and LDT accuracy (x-axis) as well as the correlation between performance on this item and WJ-Word-ID raw score (y-axis). Items that were not correlated (threshold r <= 0.10, Step 1) with overall test performance or WJ-Word-ID were removed (plotted in red). Items that passed the r>0.10 threshold are plotted in blue. Pseudowords are depicted in darker colors and real words are depicted in lighter colors. (Right panel) Difficulty (x-axis) and discriminability (y-axis, based on the 2 parameter logistic model) for each item that survived the correlation threshold. Red items were those that were removed in step 2 or 3, based on poor fit statistics of the Rasch (Step 2) or 2 parameter logistic (Step 3) models.

### Step 2: Remove words that are not well fit by the Rasch model

The Rasch model (1 parameter logistic with a guess rate fixed at 0.5) ^56^ was fit to the response data for the 429 items that remained after *Step 1* for all 118 subjects, using the MIRT package in R ^57^. Items were removed if the infit or outfit statistics were outside the range of 0.6 to 1.4, a relatively lenient criterion intended to remove items with unpredictable responses ^58^. The model was iteratively refitted until all items met this criterion. The process resulted in the removal of 129 items with response data that did not fit well with the predictions of the Rasch model (**Figure 2 right panel**, red items).

### Step 3: Examining item discrimination with the two parameter logistic model

Next we fit the two parameter logistic (2PL) model (guess rate fixed at 0.5) to the response data, for the 300 items that were retained after *Step 2*. The 2PL model fits item response functions with a difficulty parameter (*b*) as well as a slope parameter (*a*); we sought to remove any remaining items that were inefficient at discriminating different levels of reading ability based on the slope of the item response function. The right panel of **Figure 2** shows the item difficulty and discrimination parameters from the 2PL model. We removed 1 item with a low slope (*a* <0.7). Of the remaining 299 items, 114 were real words and 185 were pseudowords.

To better understand the lexical and orthographic properties of the items that were selected for the final test, we compared the retained-versus rejected-items based on word length and log frequency of their orthographic neighbors (see **Supplementary Figure 2**). We found a significant effect for word length (Kruskall-Wallis H(3) = 26.6, *P* < 0.0001): Retained real words were significantly longer than rejected real words (p <0.0001) and retained pseudowords (p = 0.050). We expected that the log frequency of the orthographic neighbors would align with the word length effect reported above (i.e,. the final set of retained words would have less frequent orthographic neighbors compared to the rejected words). Our analysis confirmed this expectation (n = 359, Kruskall-Wallis H(3)=14.5, *P*=0.0023). Indeed, pairwise comparisons confirmed that retained real words had significantly less frequent orthographic neighbors compared to the rejected real words (p <0.0004). Comparisons between retained and rejected items for bigram frequency, trigram frequency and number of morphologically complex words did not lead to any significant difference (all ps <0.05). Therefore, real words that are longer and have less frequent orthographic neighbors are retained as a result of the IRT analysis indicating their utility for measuring reading ability.

### Step 4: Test optimization

Finally, we sought to create an efficient and reliable test that was composed of three short word-lists with the following properties: (a) quick to administer through the web-browser, (b) balanced in number of real words and pseudowords, (c) matched in terms of difficulty and (d) optimally informative across the range of reading abilities. Based on feedback from subjects in *Study 1*, we surmised that a list of about 80 items was a suitable length for maintaining focus. Thus, out of the 299 items that remained after Steps 1-3, we constructed three lists with items that were equated in difficulty based on *b* parameters (estimated with the Rasch model). We further ensured that: (a) items spanned the full difficulty range to maximize test information for the lowest and highest performing participants and (b) real and pseudowords were matched in terms of length on each list. Since only 114 real words were retained after the optimization procedure described above, 76-item lists were generated with 38 pseudowords and 38 real words.

**Figure 3** shows the test information function for each of the three lists. The composite of the three word lists (234 item; scores combined across lists) was extremely reliable with a reliability of 0.97. The composite LDT has a reliability comparable to those of the WJ and TOWRE composite indices: WJ Basic Reading Skills, average reliability = 0.95 ^37^, and TOWRE Total Word Reading Efficiency, average reliability = 0.96 ^59^. Performance on the individual word lists was also highly reliable, with an ICC of 0.95 (**Figure 3 upper right**; about the same or higher than the individual TOWRE and WJ test forms: WJ Word Attack reliability = 0.90 and WJ Letter Word Identification reliability = 0.94 ^37^; TOWRE Sight Word Efficiency reliability = 0.91 and TOWRE Phonemic Decoding Efficiency reliability = 0.92 ^59^). Performance on each individual word list was highly correlated with WJ-Word-ID scores (**Figure 3 bottom panel**; List A: r = 0.88; List B: r = 0.85; List C: r = 0.87) suggesting that administering a single 80-item lexical decision task (~2-3 minutes) would be a suitable measure of reading ability for some research purposes.

**Figure 3.**
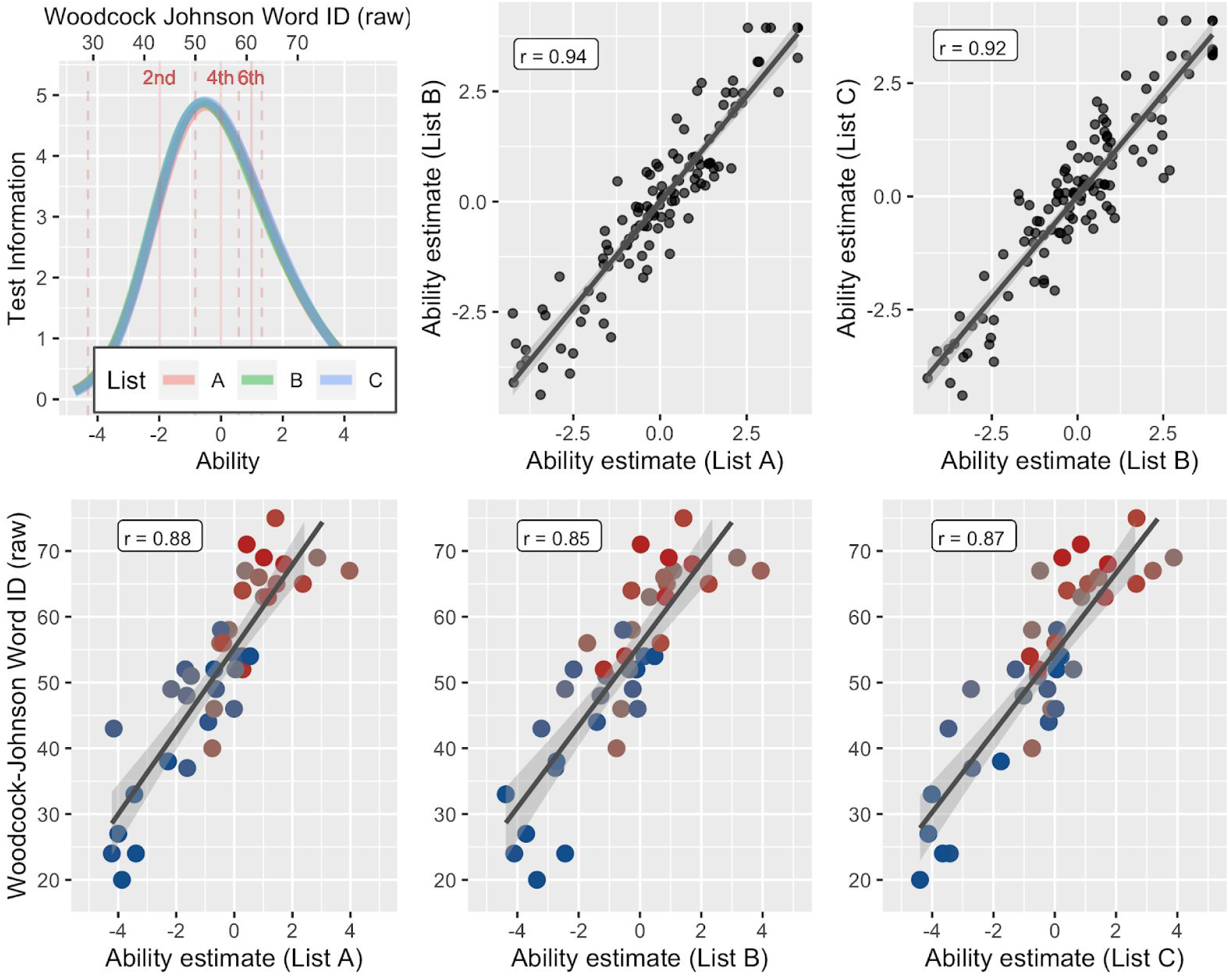
Optimized stimulus lists. (Top left) Test information functions for the three stimulus lists. The curves are overlapping over the full range, predicting near identical performance on each list irrespective of individual ability. The x-axis shows ability estimates based on the Rasch model. The upper x-axis shows the estimated WJ-Word-ID raw score equivalent based on the linear relationship between ability estimates and WJ-Word-ID scores. The pink lines indicated grade level equivalents for WJ-Word-ID scores (based on the WJ manual). Test information is high for participants scoring above the 1st grade level on the WJ-Word-ID. (Top right) Scatter plots of ability estimates based on each list. Test-retest reliability for each individual list is greater than r=0.92. (Bottom panel) Correlations between ability estimates on each list and WJ-Word-ID scores. The blue-gray-red colormap indicates standard scores as in Figure 2. Even a single short list provides an accurate and reliable estimate of WJ-Word-ID ability.

## RESULTS: STUDY 2

### Validation of optimized lexical decision task in young children

We created a new version of the browser-based LDT (*version 2*) based on the optimized word lists from *Study 1*. To improve precision in young children, twelve new words were selected from grade level text (1st and 2nd grade) and pseudoword pairs were created for each new real word (see Methods and **Supplementary Table 1** for details). 24 children between the ages of 5y11m and 7y10m (K - 2nd grade) were recruited to test whether the browser-based LDT was an accurate measure of reading ability in young children. Each child completed *version 2* of the LDT and the WJ-Word-ID was administered to each child over Zoom. All children were able to follow the instructions (which are narrated by a voice actor) and complete the task in their web-browser without assistance from a parent or researcher. Performance on the LDT was highly correlated with WJ-Word-ID in this young sample (r = 0.96, **Figure 4**).

**Figure 4.**
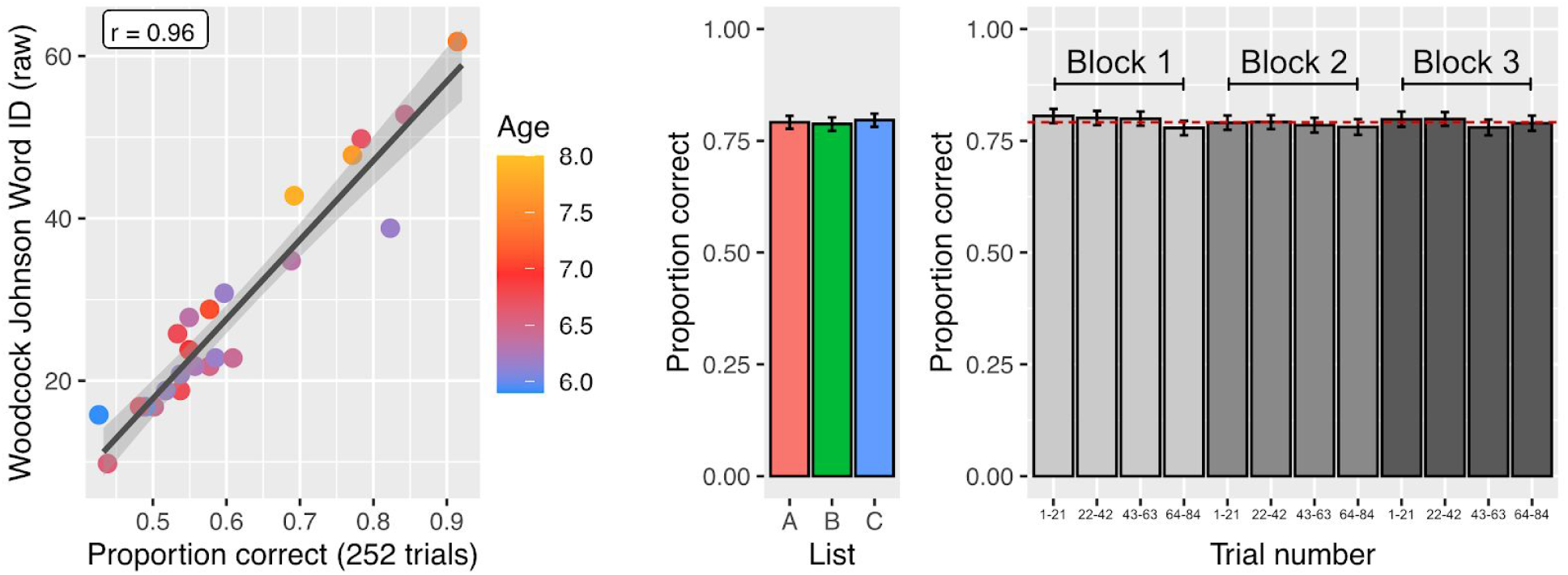
Validation of optimized lexical decision task. (left) Response accuracy on version 2 of the LDT was highly correlated with reading ability in young children. The color bar indicates the age of each child (n=24). (middle) Performance across the three matched stimulus lists was nearly identical (error bars represent +/- 1 standard error of the mean across subjects, N=118). (right) Performance is stable over the three blocks of the experiment. Each bar represents the proportion correct on 25% of the trials in a block (21 trials) and the dotted red line represents overall mean performance (N=118).

### Validation of optimized lexical decision task in children and adults

After confirming that young children could complete the browser-based LDT, we collected data on a broad age range (N = 116; 5y11m - 42y7m) and administered the WJ-Word-ID to 84 of these subjects. As expected based on *Study 1*, the optimized LDT was highly correlated with WJ-Word-ID scores (r = 0.90). Moreover, the measure was highly reliable with an intraclass correlation ^60^ of 0.91 for individual lists and 0.97 for the composite of the three lists.

### Examination of stimulus list and block order effects

The *version 2* stimulus lists were designed to be used interchangeably. **Figure 4** shows performance on the three stimulus lists (N=118); a mixed effects model confirmed that accuracy (i.e., number correct) did not differ significantly across the three lists (List B vs. List A: **β** = −0.33, SE = 0.52, t = −0.63, p = 0.53; List C vs. List A: **β** = 0.40, SE = 0.52, t = 0.77, p = 0.44). Moreover, the probability of a correct response did not change over the 252 trials (**β** = −0.013, SE = 0.02, z = −0.64, p = 0.52) and proportion correct did not differ significantly between the first, second or third block of trials (Block 2 vs. Block 1: **β** = −0.76, SE = 0.52, t = −1.47, p = 0.14; Block 3 vs. Block 1: **β** = −0.39, SE = 0.52, t = −0.75, p = 0.45).

## DISCUSSION

Our primary goal was to evaluate the suitability of a browser-based lexical decision task as a measure of reading ability. Lexical decision is commonly used to interrogate the mechanisms of word recognition, and previous studies have shown: (a) differences in task performance in dyslexia, (b) changes in task performance over development and (c) relationships to various measures of reading ability. Thus, lexical decision was a strong candidate for an automated measure of reading ability. Results from *Study 1* revealed that lexical-decision accuracy was a very accurate predictor of reading ability as measured by the WJ-Word-ID (disattenuated r = 0.94). Moreover, reliability analysis confirmed that the lexical decision task has excellent psychometric properties. Thus, by selecting the optimal list of words to span the continuum of reading ability, it might be possible to design a quick, valid and reliable measure of reading ability that could be administered to readers of all ages through the web-browser. To this end, we used IRT to select individual words with suitable measurement properties and created three short versions of the LDT that were matched in terms of difficulty. We validated this optimized version of the LDT (*version 2*) in a second study with an independent sample (*Study 2*) and confirmed the reliability and validity of this measure in children as young as six years of age. In the future, leveraging the IRT analysis to implement the LDT as a computer adaptive test would likely lead to an even more efficient and accurate measure of reading ability.

Through the process of optimizing the items to be included in the *version 2* short-form test, we made four interesting observations that are worth noting (and all the data are [publicly available](https://github.com/yeatmanlab/ROAR-LDT) for other researchers who may be interested in further item-level analyses). First, accuracy is a much better predictor of reading ability than RT, at least for children learning to read in English. Previous studies using RT have found statistically significant, but relatively weak relationships between RT and reading ability ^47^, a finding we replicate here (Results, page 4). RT is likely a useful measure for participants at the high end of the reading continuum and for children learning to read in transparent orthographies. Indeed, it is well established that accuracy quickly reaches a ceiling in transparent orthographies and individual differences are more pronounced in timed measures ^61,62^. Moreover, with a larger sample, RT might be useful for making more fine-grained discriminations of automaticity among skilled readers. Second, pseudowords are better at discriminating different levels of reading ability than are real words (as indicated by higher correlation with reading ability and more pseudowords being retained in the IRT analysis). This might reflect greater between-subject variability in performance on pseudowords compared to real words (**Supplementary Figure 2**). But it is important to keep in mind that properties of the real words included in a LDT influence responses to pseudowords and vice versa ^52,63,64^. Third, we investigated the orthographic properties of the items that were selected by the IRT analysis. Retained real words were longer and had lower frequency of orthographic neighbors than removed real words, an effect that was not observed for pseudowords (see **Supplementary Figure 1**). Finally, including items that span a wide range of difficulty (**Figure 2, 3**) is critical to a lexical-decision based measure of reading ability.

In summary, we have demonstrated that a browser-based lexical decision task is an efficient, accurate, and reliable measure of reading ability, producing metrics that are highly consistent with results of standardized, in-person reading assessment. Selection of stimuli spanning a broad difficulty range is important for the psychometric properties of the LDT. We make the Rapid Online Assessment of Reading (ROAR) openly available for other researchers studying reading development (https://github.com/yeatmanlab/ROAR-LDT-Public).

## METHODS

### Browser-based lexical decision task (version 1)

Data and analysis code are available at: https://github.com/yeatmanlab/ROAR-LDT-Public. A visual-presentation 2AFC lexical decision task (LDT) was implemented in the Psychopy experiment builder ^34^, converted to Javascript, and uploaded to Pavlovia, an online experiment platform ^65^.

The LDT was split into five blocks, each consisting of 100 trials (50 real words, 50 pseudowords) and each block was introduced as a quest in the magical world of Lexicality. To engage a wide range of subjects, the LDT was embedded in a fun story, animated environment, and a game that involved collecting golden coins. Instructions were narrated by a character in the game and there was a series of 10 practice trials with feedback to ensure that each participant understood the LDT. After a practice trial with an incorrect response, the character would return, explain why the response was incorrect, and remind the participant the rules of the LDT. This ensured that even the youngest participants understood the rules of the LDT and remained engaged through the duration of the task.

Prior to the five blocks of the LDT, subjects completed a simple response time (RT) task. This task was used as a baseline measure, in case there was too much noise in the response time data (e.g., due to different devices being used). In each trial of the simple RT task, participants saw a fixation cross, then a triangle or a square flashed briefly for 350 ms, followed by another fixation cross. Participants were instructed to use the keyboard to respond, by pressing “left arrow” to indicate that a triangle was presented, or “right arrow” to indicate that a square was presented.

The five blocks of the LDT followed a similar format: a word or pseudoword (Arial font) would flash briefly for 350 ms in the center of the screen, followed by a fixation cross. Participants pressed “right” for a real word or “left” for a pseudoword. Stimuli were presented in the center of the screen and scaled to be 3.5% of the participants’ vertical screen dimension.

The task was gamified in order to be more engaging for the younger participants. If the participants achieved ten consecutive correct answers, a small, one-second animation would play showing their party of adventurers defeating a group of monsters, with a small “+10 gold” icon appearing on top. After each block, participants unlocked new characters and were shown the amount of gold that they had collected during the last block.

### Word and Pseudoword Stimuli (version 1)

250 real words and 250 pseudowords were selected to span a large range of lexical and orthographic properties, with the goal of finding stimuli spanning a large range of difficulty. The distribution of lexical and orthographic properties of the stimuli is shown in **Supplementary Figure 1**. Pseudowords were matched in orthographic properties to real words using Wuggy, a pseudoword generator ^66^. Wuggy generates orthographically plausible pseudowords that are matched in terms of sub-syllabic segments, word length, and letter-transition frequencies. Matched real word/pseudoword pairs were kept within the same block, such that each block contained 50 real words and 50 pseudowords roughly matched in terms of orthographic properties and stimulus order was randomized. Additionally, 18 real words were paired with 18 orthographically implausible pseudowords to create some easier items to ensure that the LDT would not have floor effects for young children and those with dyslexia (e.g. “wndo”). These orthographically implausible words had low trigram frequencies and can be appreciated from the bump at the lower end of the trigram frequency distribution in the lower panel of **Supplementary Figure 1**.

### Participant recruitment and consent procedure (Study 1)

Each adult participant provided informed consent under a protocol that was approved by the Institutional Review Board at Stanford University or The University of Washington and all methods were carried out in accordance with these guidelines. Each participant under 18 years of age provided assent and a parent or legal guardian provided informed consent.

Participants were recruited from two research participant databases: (1) The Stanford University Reading & Dyslexia Research Program (http://dyslexia.stanford.edu) and (2) The University of Washington Reading & Dyslexia Research Program (http://ReadingAndDyslexia.com). Each database includes children and adults who have (1) enrolled and consented (and/or assented) to being part of a research participant pool, (2) filled out extensive questionnaires on demographics, education history, attitudes towards reading and history/diagnoses of learning disabilities, (3) been validated through a phone screening to ensure accuracy of basic demographic details entered in the database. Many of the children and adults have participated in one or more study at Stanford University or the University of Washington and, as part of these studies, have undergone an in-person assessment session including standardized measures of reading abilities (Woodcock-Johnson IV Tests of Achievement, Test of Word Reading Efficiency - 2), verbal abilities (Welschler Abbreviated Scales of Intelligence II (WASI-II), Vocabulary subtest) and general reasoning abilities (WASI-II, Matrix Reasoning subtest). Scores on standardized tests were analyzed if the testing had occurred within 12 months of completing the LDT task (scores were adjusted based on the expected improvement since the time of testing). A WASI IQ score less than 70 has been used as an exclusion criteria in previous studies with this subject pool and, therefore, subjects with low IQ were excluded from the present study (when scores were available).

Participants in the database were emailed a brief description of the study and a link to a digital consent form and the online LDT task. If participants provided consent, their LDT performance was linked to their assessment data in the research database (questionnaires and standardized tests). Participants also had the option to complete the online experiment and remain anonymous. A total of 120 children and adults between the ages of 6.27 years and 29.15 (M = 11.61, SD = 3.67) participated in *Study 1*. These subjects spanned the full range of reading abilities as measured by the Woodcock Johnson Basic Reading Skills (WJ-BRS) standard scores (M = 97.7, SD = 18.1, min = 57, max= 142) and TOWRE Index standard scores (M = 92.4, SD = 18.9, min = 55, max= 134). Many subjects were recruited for studies on dyslexia and, based on the WJ-BRS, 25% of the subjects met a typical criterion for dyslexia (standard score more than 1 SD below the population mean). 37.5% of subjects were below the 1 SD cutoff in terms of TOWRE Index. We treat reading ability as a continuous measure throughout as opposed to imposing a threshold that defines a group of participants as individuals with dyslexia. Demographic information for the participants can be found at:https://github.com/yeatmanlab/ROAR-LDT-Public/blob/main/data_allsubs/demographic_all_newcodes.csv

### Data analysis (Study 1)

All analysis code and data is publicly available at: https://github.com/yeatmanlab/ROAR-LDT-Public, with a README file that documents how to reproduce each figure and statistic reported in the manuscript. Correct/incorrect responses and log transformed response times (RTs) for each item were concatenated across subjects into a large table. Two subjects were identified as outliers and removed from further analysis based on the following procedure: First, median RTs were calculated for each participant. Then, participants were excluded if their median RT was more than 3 standard deviations below the sample mean. The two subjects who met this criterion performed the LDT at near chance accuracy level, suggesting that their extremely fast RTs were indicative of random guessing (lack of experiment compliance). These participants are displayed in the figures as open circles for reference. For the analysis of RT data, responses shorter than 0.2 seconds or longer than 5 seconds were removed. Then, quartiles and the interquartile range (IQR) of the RT distribution were calculated per participant. Responses that were longer than 3 times the IQR from the third quartile, or shorter than 3 times the IQR from the first quartile, were further excluded (see ^67,68^ for similar outlier exclusion procedures). These steps resulted in the exclusion of 4.6% of total trials. Analysis of RT data was conducted using the lme4 package in R ^69^.

### Browser-based lexical decision task (version 2)

A shorter version of the browser-based LDT (252 trials in *Study 2* vs. 500 trials in *Study 1*) was implemented based on the three, optimized 76-item lists that were designed in *Study 1*. Each list is presented in a block of trials and the order of the lists is randomized as are the order of the stimuli within each list (but stimuli were not mixed across lists). Otherwise, all the details of the experiment were maintained from *version 1*.

Twelve new words were selected from grade level text (1st and 2nd grade) in order to include easy vocabulary words for young elementary school children (and English language learners). Pseudoword pairs were created for each new real word by rearranging the letters and maintaining orthographic regularities (as in *version 1*). These new items (4 real words, 4 pseudowords) were added to each list for a final length of 84 items per list. The three stimulus lists for *Study 2* are shown in **Supplementary Table 1**.

### Participant recruitment (Study 2)

N=116 children and adults between the ages of 5y11m - 42y7m were recruited through: (1) partnerships with local school districts, (2) research participant databases (http://dyslexia.stanford.edu), and (3) ongoing studies. One of the goals of *Study 2* was to validate the Rapid Online Assessment of Reading (ROAR) LDT in young children (< 8 years of age). Thus, a specific effort was made to recruit participants who had just finished kindergarten (n = 20). Each participant who completed the ROAR was invited to participate in an assessment session conducted over Zoom in which we administered the Woodcock Johnson Word ID (n = 84). These data were used to confirm the results of *Study 1*, and validate *version 2* of the ROAR in an independent sample.

## ACKNOWLEDGEMENTS

We would like to thank the Pavlovia and PsychoPy team for their support on the browser-based experiments. This work was funded by NIH NICHD R01HD09586101, research grants from Microsoft and Jacobs Foundation Research Fellowship to J.D.Y.

## AUTHOR CONTRIBUTIONS

JDY and KAT designed the study. KAT, PMD and MET collected the data. JDY, KAT, PMD, MY, MR, IIK, SC, KK, MB and BWD analyzed the data. All authors contributed to the writing of the manuscript.

## SUPPLEMENTARY MATERIAL

**Supplementary Figure 1.**
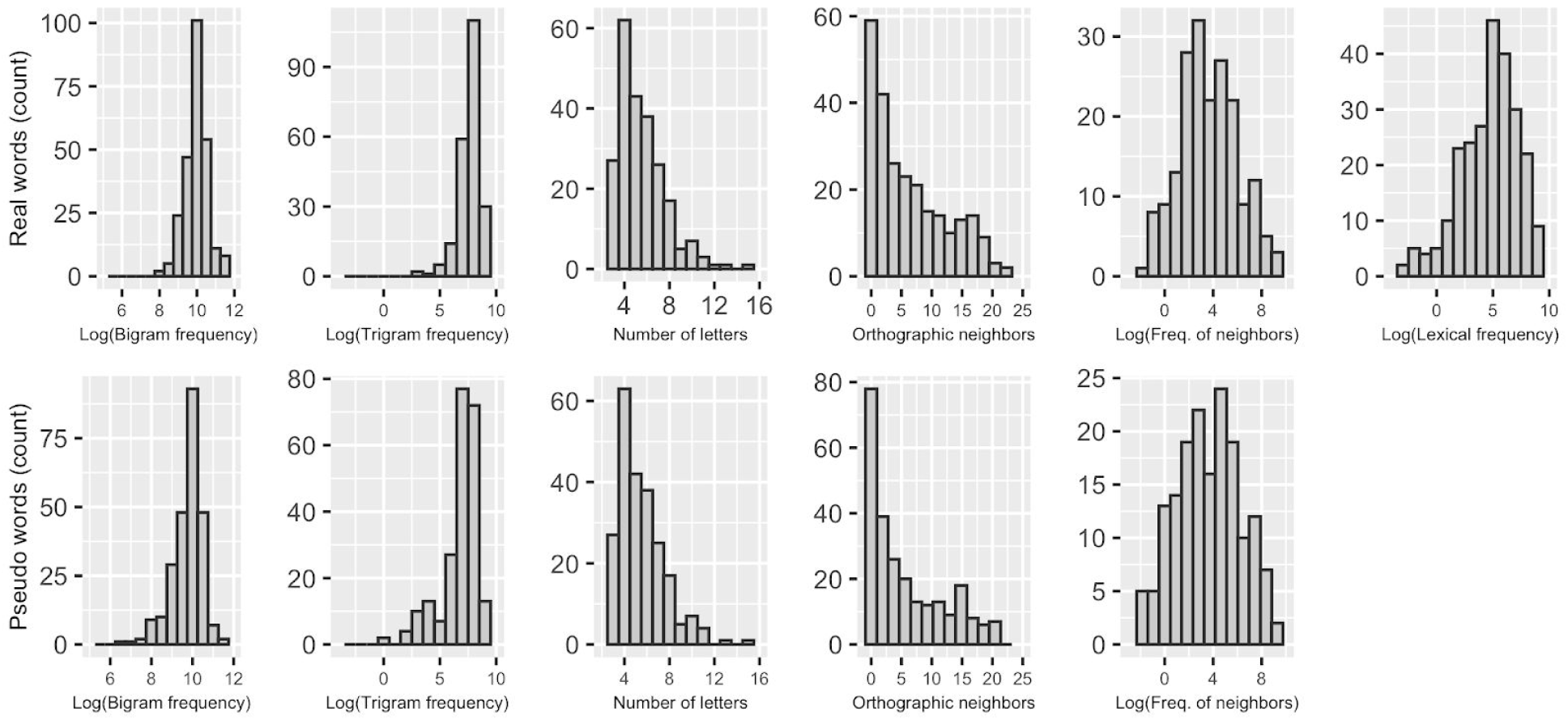
Lexical and orthographic properties of the stimuli. (Top panel) Bigram frequency, trigram frequency, number of letters, number of orthographic neighbors, frequency of orthographic neighbors and lexical frequency for the 250 real words used in the lexical decision task (LDT). (Bottom panel) Bigram frequency, trigram frequency, number of letters, number of orthographic neighbors, frequency of orthographic neighbors for the 250 pseudo words used in the LDT.

**Supplementary Figure 2:**
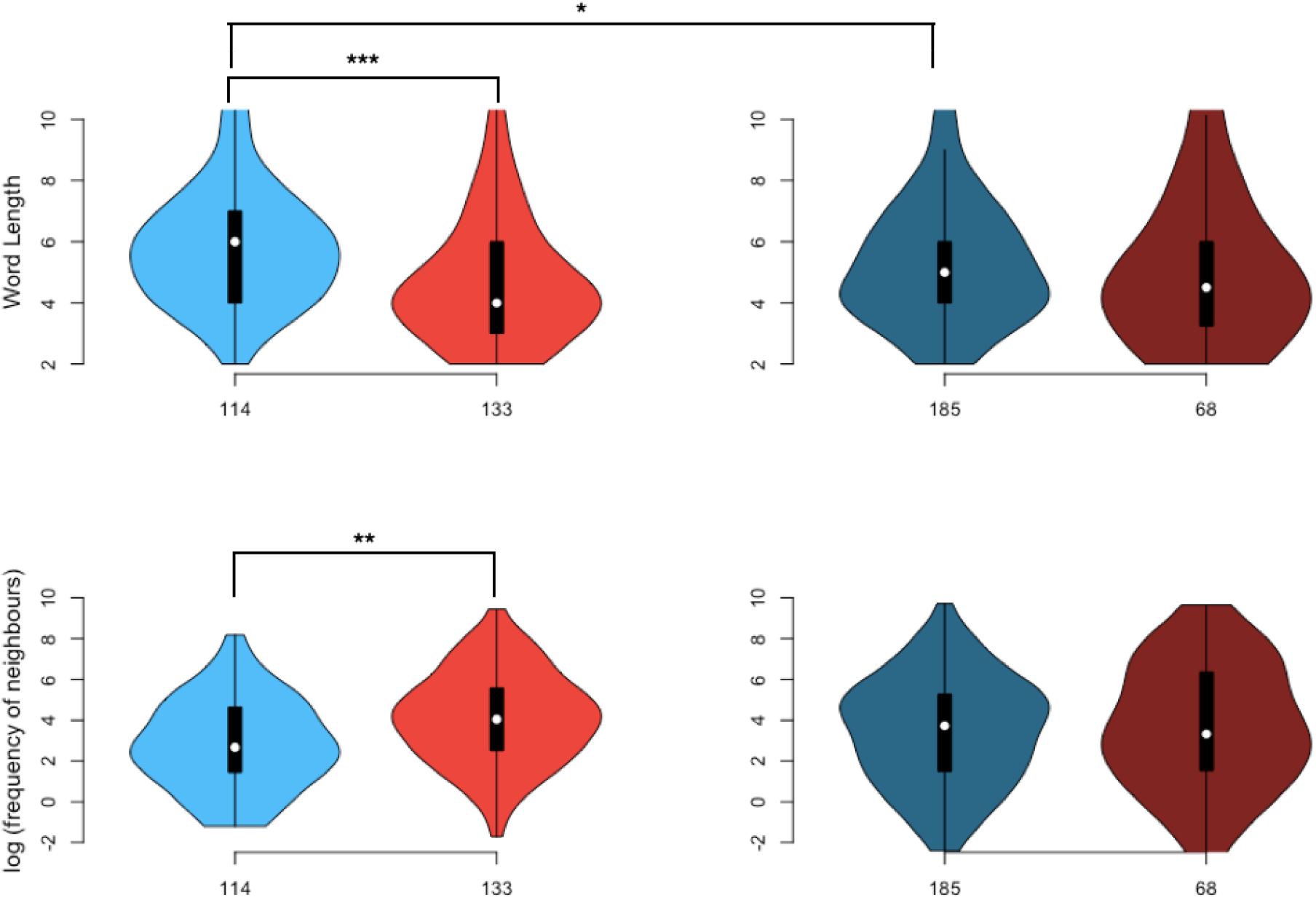
Top Panel (left: real words; right: pseudowords): shows the distribution of word length for the retained real words (BLUE) and rejected real words (RED) from step 3 of the IRT analysis. We found that the retained real words were longer than rejected real words (p <0.0001***) and retained pseudowords (p =0.050*). Bottom Panel (left: real words; right: pseudowords): Pairwise comparisons revealed that retained real words were significantly lower in log frequency of its orthographic neighbors compared to the rejected real words (p <0.0004**). These analyses show that the more unique a word is (longer and with low-frequency neighbors) the higher its discriminative power of reading performances. Darker colors are pseudowords and lighter colors are real words.

## Optimized Stimulus Lists (ROAR *Version 2*)

**Supplementary Table 1.**
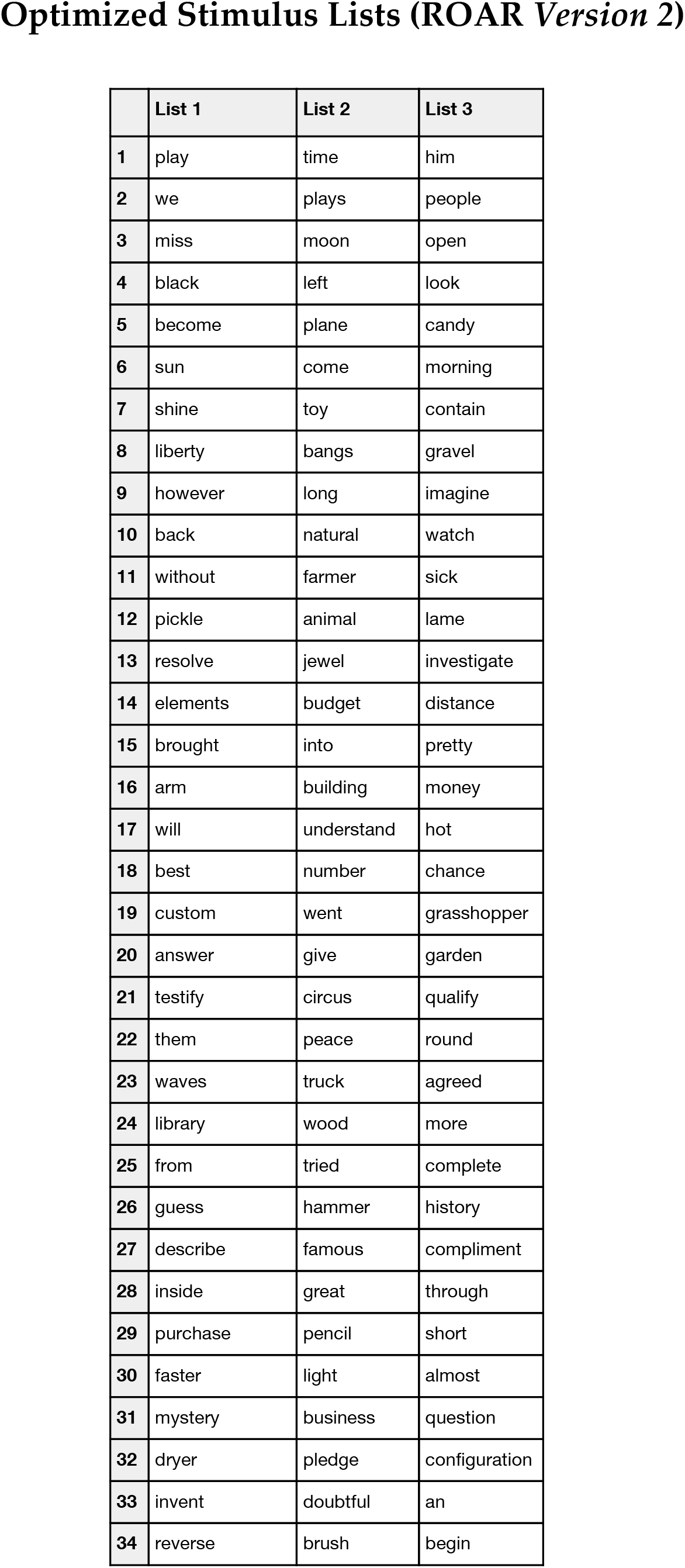
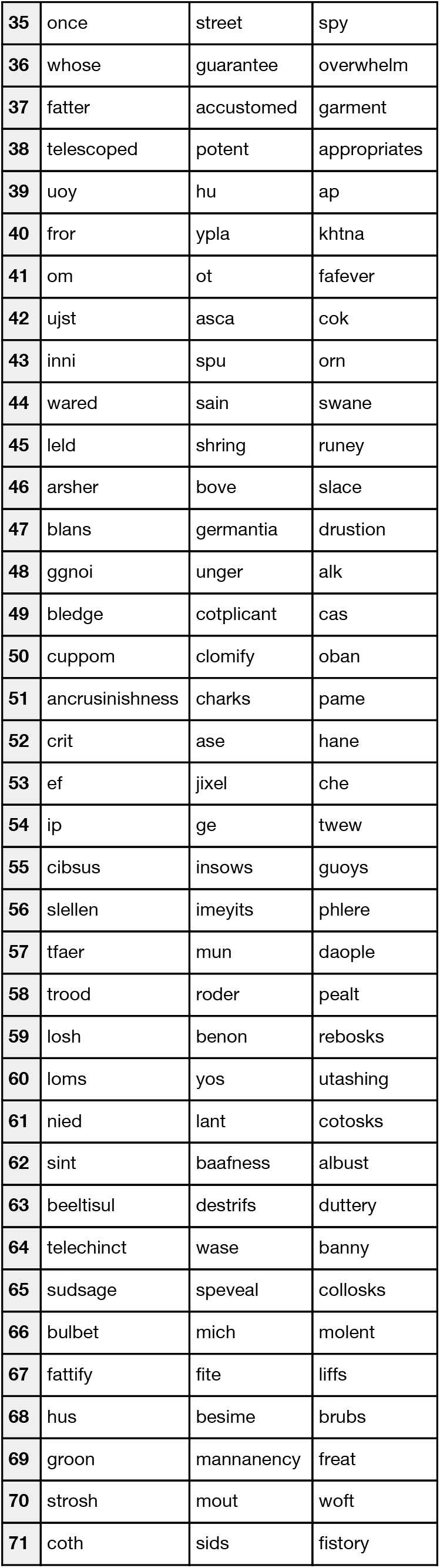
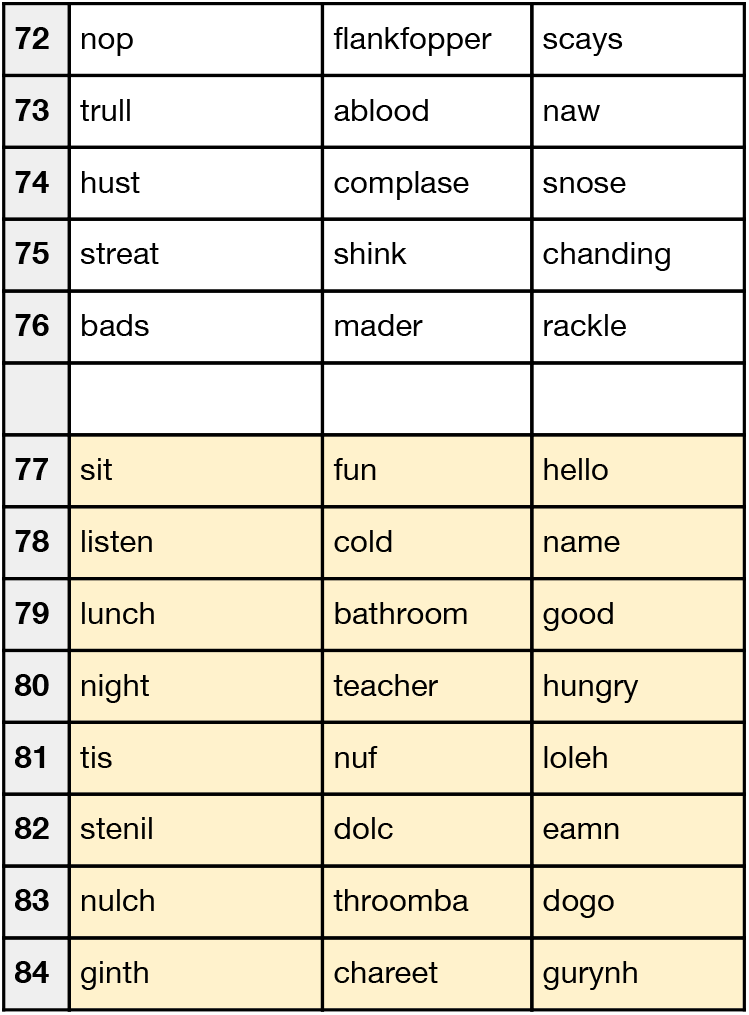
Stimuli for ROAR version 2 used in Study 2. 76 items (38 real words, 38 pseudowords) were selected based on the data from Study 1. Eight new items (4 real words, 4 pseudowords) were added to each list to include more easy vocabulary words for young children and English language learners. Yellow cells at the bottom of each list indicate the new items. Final stimulus lists were 84 items long.

**Supplementary Table 2.** Percent correct and response times for real and pseudo words. Mean and standard deviation is shown for response time (RT) and percent correct

|  | Mean RT | SD RT | Mean % correct | SD % correct |
| --- | --- | --- | --- | --- |
| Pseudo Word | 0.934 | 0.266 | 74.3 | 16.4 |
| Real Word | 0.831 | 0.220 | 79.8 | 12.8 |

